# A Single Multipurpose FSH–Blocking Therapeutic for Osteoporosis and Obesity

**DOI:** 10.1101/2022.02.28.482279

**Authors:** Sakshi Gera, Tan-Chun Kuo, Funda Korkmaz, Damini Sant, Victoria DeMambro, Anisa Gumerova, Kathayani Sudha, Ashley Padilla, Geoffrey Prevot, Jazz Munitz, Abraham Teunissen, Mandy van Leent, Tomas G.J.M. Post, Jessica C. Fernandes, Jessica Netto, Farhath Sultana, Eleanor Shelly, Pushkar Kumar, Liam Cullen, Jiya Chatterjee, Sari Miyashita, Hasni Kannangara, Megha Bhongade, Kseniia Ievleva, Valeriia Muradova, Rogerio Batista, Cemre Robinson, Anne Macdonald, Susan Babunovic, Mansi Saxena, Marcia Meseck, John Caminis, Jameel Iqbal, Maria I. New, Vitaly Ryu, Se-Min Kim, Jay J. Cao, Neeha Zaidi, Zahi Fayad, Daria Lizneva, Clifford J. Rosen, Tony Yuen, Mone Zaidi

## Abstract

Pharmacological and genetic studies over the past decade have established FSH as an actionable target for diseases affecting millions, notably osteoporosis, obesity and Alzheimer’s disease (AD). Blocking FSH action prevents bone loss, fat gain and AD–like features in mice. We recently developed a first–in–class, humanized, epitope–specific FSH blocking antibody, MS-Hu6, with a K_D_ of 7.52 nM. Using a GLP–compliant platform, we now report the efficacy of MS-Hu6 in preventing obesity and osteoporosis in mice, and parameters of acute safety in monkeys. Biodistribution studies using ^89^Zr–labelled, biotinylated or unconjugated MS-Hu6 in mice and monkeys showed localization to bone, bone marrow and fat depots. MS-Hu6 displayed a β phase t_½_ of 13 days (316 hours) in humanized *Tg32* mice, and bound endogenous FSH. We tested 215 variations of excipients using the protein thermal shift assay to generate a final formulation that rendered MS-Hu6 stable in solution upon freeze–thaw and at different temperatures, with minimal aggregation, and without self–, cross–, or hydrophobic interactions or appreciable binding to relevant human antigens. MS-Hu6 showed the same level of “humanness” as human IgG1 *in silico*, and was non–immunogenic in ELISPOT assays for IL-2 and IFNγ in human peripheral blood mononuclear cell cultures. We conclude that MS-Hu6 is efficacious, durable and manufacturable, and is therefore poised for future human testing as a multipurpose therapeutic.

## INTRODUCTION

While obesity and osteoporosis are both diseases of public health concern, the paucity of therapies to prevent and treat them continues to represent a challenge (1, 2). Accumulating clinical data suggest that the two disorders track together in women across the menopausal transition. Particularly during the late perimenopause, there is precipitous bone loss, onset of visceral obesity, dysregulated energy balance, and reduced physical activity (3–9). These aberrant physiologic changes across the menopausal transition are not fully explained by low estrogen, as estrogen levels are relatively unperturbed, while serum FSH levels rise to maintain estrogen secretion from an otherwise failing ovary (10–12).

The question has been whether a rising FSH level is a driver for post–menopausal obesity and osteoporosis. In 2006, we provided the first evidence for a direct action of FSH on bone (13). Since then, despite controversy fueled mainly from the over–interpretation of clinical studies with GnRH agonists that suppress not only FSH, but also GnRH and LH (14), there is replicable evidence that the selective inhibition of FSH action in mice, for example by using novel FSH– blocking antibodies or a GST–FSH fusion protein as a vaccine, protects against hypogonadal bone loss (15–19). Serum FSH, bone turnover, and bone mineral density also correlate well in women, particularly when FSH levels are rising during the late perimenopause [review: (20)]. Likewise, activating *FSHR* polymorphisms in postmenopausal women are linked to a high bone turnover and reduced BMD (21). Thus, it makes both biological and clinical sense to selectively inhibit FSH action to prevent bone loss.

We and our collaborators have also shown that inhibiting FSH by FSH–blocking antibodies reduces white adipose tissue (WAT) in every fat compartment, induces thermogenic (or beige) adipose tissue, and increases energy expenditure in mice (16). Reduced fat mass has also been documented with a vaccine containing tandem repeats of the 13–amino–acid–long FSH receptor–binding FSHβ sequence to which our antibodies were raised (22). An interventional study in treatment–naïve prostate cancer patients comparing orchiectomy *versus* triptorelin showed that, with near–zero testosterone, patients on triptorelin (reduced serum FSH and LH) had significantly lower body weight and fat mass compared to those post–orchiectomy (23). Even recognizing the constraints of using a GnRH agonist, this dataset suggests that lowering serum FSH could, in principle, have beneficial effects on body composition in people, despite concomitant reductions in GnRH and LH. There is also new evidence that selective FSH blockade lowers serum total and LDL cholesterol (24, 25).

We hypothesize that blocking FSH action will reduce obesity and bone loss in people. Towards this goal, we have developed our lead candidate, a first–in–class humanized FSH– blocking antibody, MS-Hu6. The latter binds a 13–amino–acid–long epitope of human FSHβ (LVYKDPARPKIQK) with high affinity, and by doing so, blocks the interaction of FSH with its receptor (26). Here, we report a comprehensive characterization of MS-Hu6 in terms of its *in vivo* efficacy in mouse models of obesity and osteoporosis, acute safety in monkeys, a full evaluation of its pharmacokinetic, pharmacodynamic and biodistribution, and a compendium of its physicochemical properties. This new information provides the framework for first–in–human studies towards the future use of MS-Hu6 in obesity and, osteoporosis.

## RESULTS

### Efficacy of MS-Hu6 in Reducing Body Fat and Inducing Beige Adipose Tissue

In choosing MS-Hu6 as the lead candidate from an array of 30 humanized clones, we examined the electrostatic binding *in silico* and determined K_D_ by surface plasmon resonance *in vitro* (26). MS-Hu6 had the best affinity (K_D_ = 7.52 nM), approaching that of trastuzumab. We fine mapped the three top candidates to document subtle differences in binding modes (26). In addition, we established that MS-Hu6 blocked the binding of labeled recombinant human FSH to the FSHR, and in doing so, inhibited osteoclastogenesis and promoted beiging of adipocytes *in vitro* (26). Here, we studied the effect of MS-Hu6 on body composition in male ThermoMice fed *ad libitum* on a high fat diet. Mice were injected with MS-Hu6 or human IgG (7 µg/day, 5 days– a–week) for 8 weeks. The latter dose was based on the *in vitro* IC_50_ of MS-Hu6, which was ∼30– fold lower than our polyclonal Ab (26). We determined net food intake and measured body weight weekly. Of note was a trend towards an increase in food intake in MS-Hu6–treated mice compared with those given human IgG (Fig. 1A). Despite this trend, there was a decline in body weight, with statistically significant decrements at weeks 7 and 8 (Fig. 1A). We also performed quantitative nuclear magnetic resonance (qNMR) at week 8—this revealed a significant decrease in total mass and fat mass with an increase in lean mass (Fig. 1B). This data mimics FSHR haploinsufficiency in male *Fshr*^+/-^ mice (Fig. 1C). Reduced fat mass was notable on manual weighing in the mesenteric, renal, and gonadal compartments—and, expectedly not in brown adipose tissue (BAT) in MS-Hu6–injected mice (Fig. 1D) (16). Quantitative PCR showed evidence for reduced expression of fat genes, namely *Pparg*, *Fabp4* and *Cebpa*—either significantly or with trends—in subcutaneous and gonadal WAT and BAT (Fig. 1E). Of note is that serum LH, GnRH and testosterone, levels were unchanged after 8 weeks of MS-Hu6, with an unexplained drop in in serum activin (Fig. 1F).

**Figure 1:**
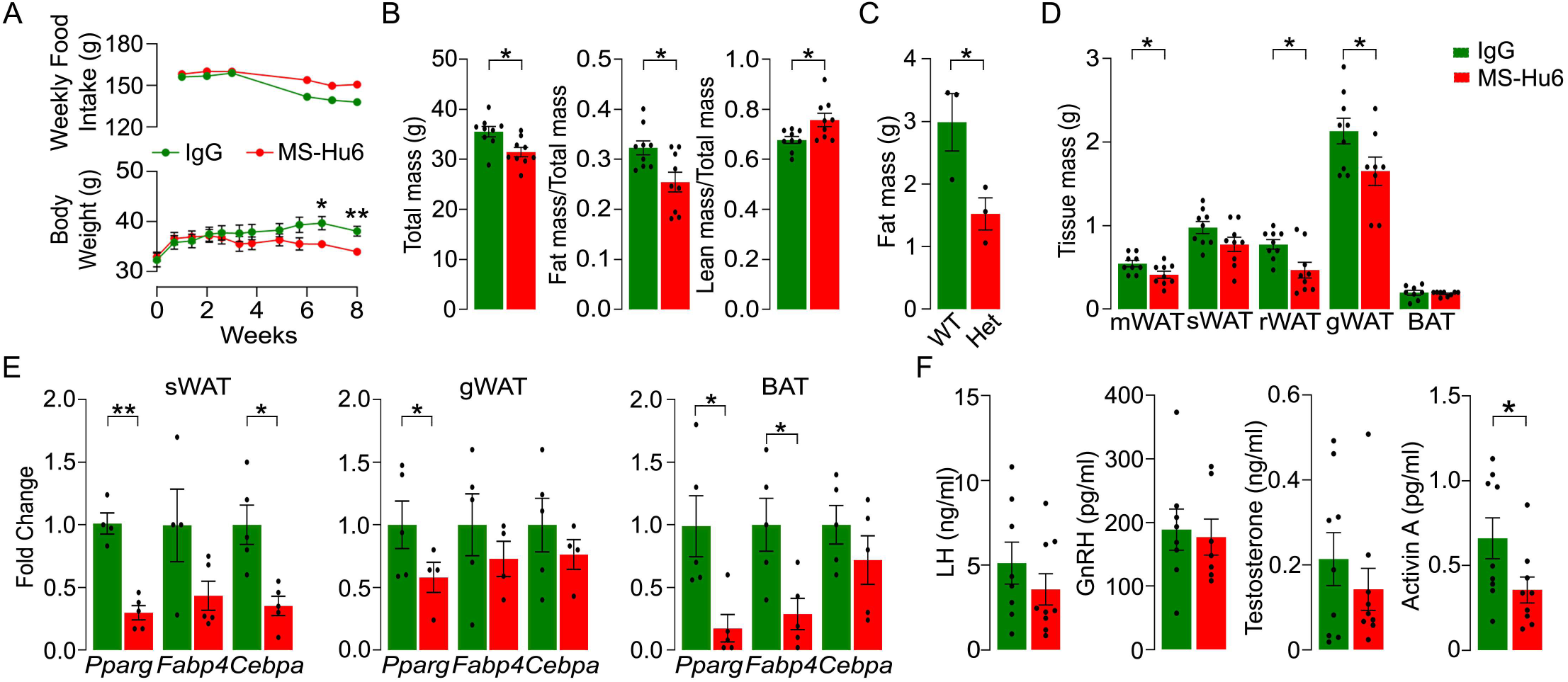
MS-Hu6 Reduces Body Weight and Body Fat. Effect of MS-Hu6 or human IgG (7 µg/day, 5 days–a–week) injected i.p. for 8 weeks on food intake and body weight **(A)**, and total mass, fat mass/total mass and lean mass/total mass (by quantitative NMR) **(B)** in male ThermoMice fed *ad libitum* on a high fat diet. The effect of MS-Hu6 in reducing body fat mimics the effect of *Fshr* haploinsufficiency in *Fshr^+/-^* mice **(C)**. Weight–based measurements of white adipose tissue in various compartments, namely mesenteric (mWAT), subcutaneous (sWAT), renal (rWAT) and gonadal (gWAT), as well as interscapular brown adipose tissue (BAT) **(D)**. Quantitative PCR showing the expression of fat genes, namely *Pparg*, *Fabp4* and/or *Cebpa* were reduced in sWAT, gWAT and/or BAT **(E)**. Serum LH, GnRH, testosterone, activin A and total inhibin levels **(F)**. Statistics: mean ± s.e.m.; *N*=9 mice/group for panels A, B and D; *N*=3 mice/group for panel C; 4 to 5 biological replicates for panel E; *N*=7-9 mice/group for panel F; \**P*<0.05, \*\**P*<0.01.

We explored the activation of interscapular BAT and beiging of WAT *in vivo* by IVIS imaging after 4 and 8 weeks of MS-Hu6 or human IgG treatment. ThermoMice harbor a *Luc2– T2A–tdTomato* dual reporter transgene on the Y chromosome driven by the *Ucp1* promoter (27) (Fig. 2A). Mice were injected with D-Luciferin (150 mg/kg), followed by imaging at 3, 5, 10 and 12 minutes to calculate average radiance. The peak radiance at 10 minutes was higher at 8 weeks, with a trend at 4 weeks in mice receiving MS-Hu6 compared with those given human IgG (Fig. 2B). The upper dorsal interscapular region showed profound BAT activation. WAT beiging and BAT activation were both confirmed by UCP1 immunohistochemistry in formalin–fixed, paraffin–embedded sections (Fig. 2C). Of note is that the UCP-1–high beige adipocytes in MS-Hu6–treated mice appeared highly condensed (Fig. 2C). This effect on cell size was confirmed by morphometry in hematoxylin/eosin stained sections (Fig. 2D). Certain beiging genes, including *Ucp1*, *Cox7* and/or *Cidea* were upregulated in mice given MS-Hu6, again consistent with white– to–beige transition (Fig. 2E).

**Figure 2:**
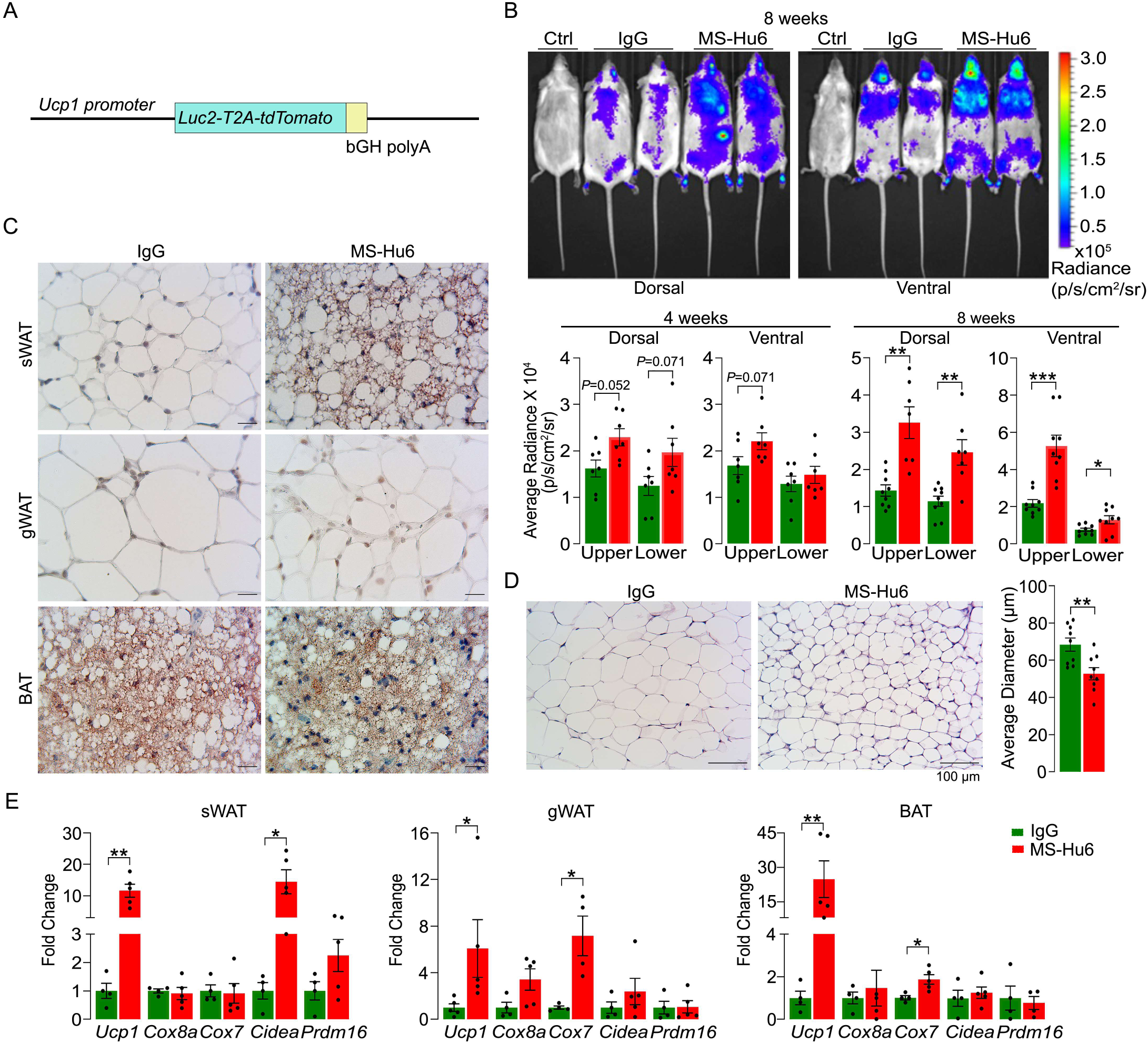
MS-Hu6 Induces Beiging of White Adipose Tissue. **(A)** Transgenic construct of *Luc2-T2A-tDtomato* under control of the *Ucp1* promoter. **(B)** Effect of MS-Hu6 or human IgG (7 µg/day, 5 days–a–week, i.p.) in male ThermoMice fed *ad libitum* on a high fat diet on emitted radiance from dorsal or ventral surfaces after 4 and 8 weeks post–injection following D-Luciferin (IVIS, 10 minute peak, average radiance, Ctrl: non–transgenic mice given D-Luciferin). **(C)** UCP1 immunohistochemistry of subcutaneous (sWAT) and gonadal white adipose tissue (gWAT) and brown adipose tissue (BAT). **(D)** Hematoxylin–eosin staining of WAT showing cell condensation in MS-Hu6–treated mice, measured morphometrically as average (Avg) diameter. **(E)** Quantitative PCR showing increased expression of beiging genes, namely *Ucp1*, *Cox8a, Cox 7, Cidea* and/or *Prdm16* in sWAT, gWAT and/or BAT in MS-Hu6–treated mice. Statistics: mean ± s.e.m.; *N*=7 and 9 mice/group for panels B and D, respectively; 4 to 5 biological replicates for panel E; \**P*<0.05, \*\**P*<0.01, \*\*\**P*<0.001, and as shown.

### Efficacy of MS-Hu6 in Improving Bone Mass

We examined the effect of MS-Hu6 on mouse bones in ThermoMice after 8 weeks of MS-Hu6 or human IgG treatment. Histomorphometry of femoral metaphysis showed significant increases in fractional bone volume (B.Ar/T.Ar) and trabecular thickness (Tb.Th), without an effect on trabecular number (Tb.N) (Fig. 3A). This increase in fractional bone volume was consistent with areal BMD measures (Fig. 3B). Dynamic histomorphometry showed evidence for increased mineral apposition rate (MAR) and bone formation rate (BAR) (Fig. 3C), confirming increased osteoblastic bone formation. In line with a predominately anabolic action of MS-Hu6, the expression of osteoblast genes in bone extracts, such as *Col1a1*, *Alp* and *Runx2*, were either significantly increased or showed trends (Fig. 3D). In contrast, while osteoclast surface (Oc.s/BS) was not reduced, there was a reduction in the expression of the osteoclast marker gene *Acp5* (Fig. 3E). The latter finding is not surprising as male ThermoMice mice were not in a high bone turnover state, in which instance a further lowering of bone resorption from baseline would not normally be expected. However, the *ex vivo* reduction of *Acp5* expression suggests that MS-Hu6 was inhibiting the effect of FSH on TRAP–positive osteoclasts.

**Figure 3:**
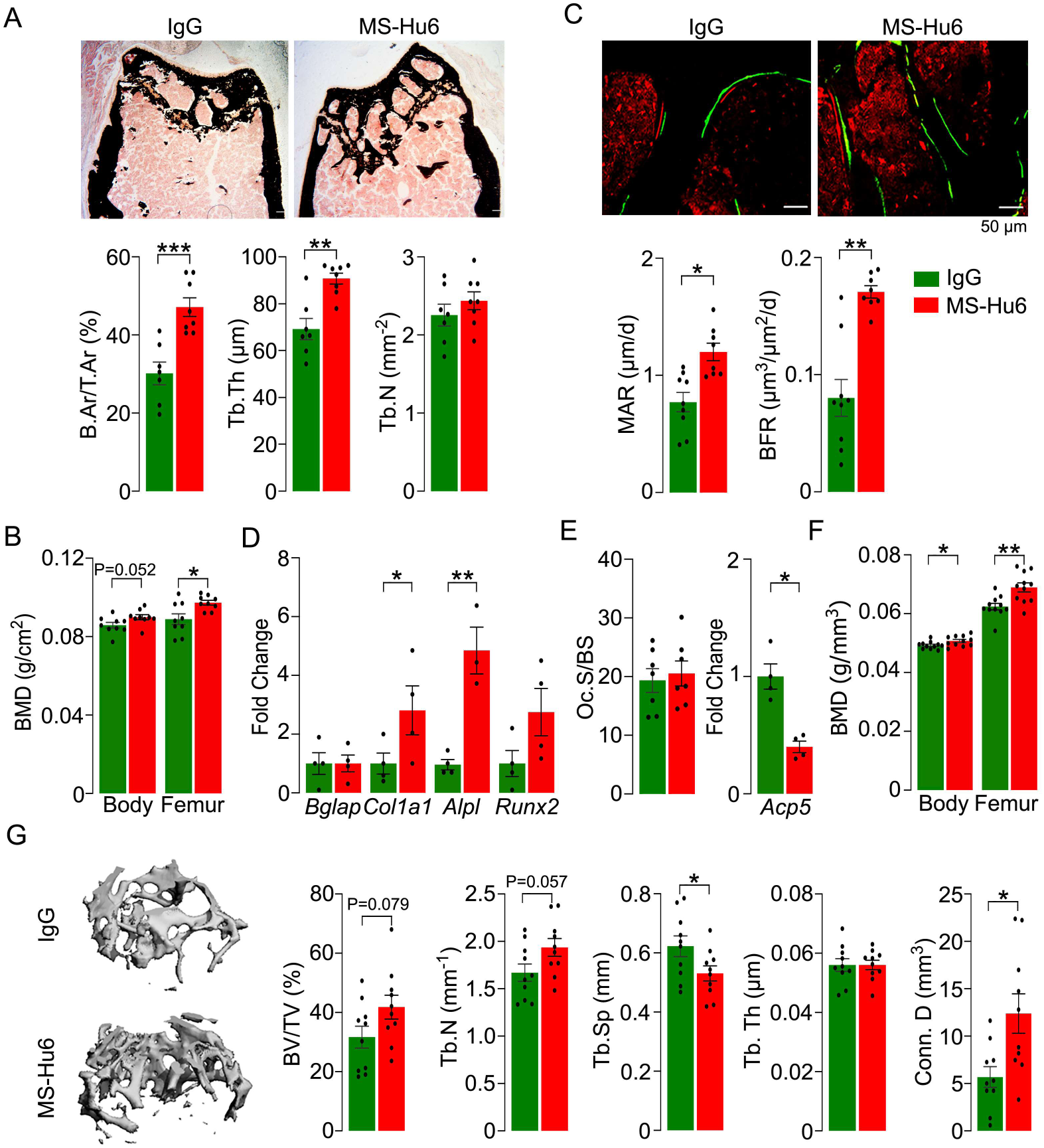
MS-Hu6 Stimulates New Bone Formation and Increases Bone Mass. **(A)** Representative images of von Kossa–stained femoral epiphyses of male ThermoMice mice that were treated with MS-Hu6 or human IgG (7 µg/day, 5 days–a–week, i.p.) and fed *ad libitum* on a high fat diet. Two–dimensional histomorphometric parameters showing bone volume (B.Ar/T.Ar), and trabecular thickness (Tb.Th) and number (Tb.N). The increase in B.Ar/T.Ar was confirmed on BMD measurements in live mice **(B)**. Dynamic histomorphometry showing representative images of double–labelled sections and quantitative data on mineral apposition rate (MAR) and bone formation rate (BFR). **(D)** Quantitative PCR showing the expression of osteoblastogenesis genes, namely *Bglap*, *Col1a1*, *Alpl* and *Runx2*. **(E)** Osteoclast surfaces (Oc.S.) remained unchanged, but with a reduction in *Acp5* (TRAP) gene expression. Parallel studies carried out at C.J.R.’s lab used 3–month–old ovariectomized C57BL/6 mice, which were injected 24 weeks post–ovariectomy with MS-Hu6 or human IgG, daily, at 100 µg/day, for 4 weeks and then 50 µg/day for 4 weeks. Shown are *Piximus* measurements of total body and femur bone mineral density (BMD) **(F),** as well as µCT images and quantitative estimates of fractional bone volume (BV/TV), trabecular number (Tb.N), spacing (Tb.S.) and thickness (Tb.Th), and connectivity density (Conn.D) (performed at J.C.’s lab). Statistics: mean ± s.e.m.; *N*=7, 8 mice/group for panel A; *N*=9 mice/group for panel B; *N*=9, 8 mice/group for panel C; 4 biological replicates for panel D; *N*=7 mice/group and 4 biological replicates for the left and right bar graphs, respectively, for panel E; *N*=11 and 10 mice/group for panel F and G, respectively; \**P*<0.05, \*\**P*<0.01, \*\**P*<0.001, and as shown.

To further explore the anabolic action of MS-Hu6, and to replicate our dataset in C.J.R.’s lab, 3–month–old female C57BL/6 mice were ovariectomized, followed 24 weeks later by injection of MS-Hu6 or human IgG, daily, at 100 µg/day, for 4 weeks and then 50 µg/day for a further 4 weeks. Total body and femoral BMD measured by *Piximus* was increased significantly in mice treated with MS-Hu6 (Fig. 3F). Micro CT of the femoral epiphysis showed increased BV/TV (*P*=0.079), Conn.D (*P*<0.05), and Tb.N (*P*=0.057), reduced Tb.Sp (*P<*0.05), and no change in Tb.Th (Fig. 3G). Expectedly, these anabolic effects were not seen in similarly treated C3H/HeJ mice, which are known to display a high bone mass phenotype (Supplementary Fig. 1) (28). Overall, therefore, our data show that consistent with previous studies using our polyclonal antibody (17), MS-Hu6 displays an anabolic action in replenishing lost bone.

### Pharmacokinetics and Pharmacodynamics of MS-Hu6

Pharmacokinetic studies were performed in three mouse models—C57BL/6, CD1 and *Tg32* mice—using ^89^Zr–labelled, unconjugated, or biotinylated MS-Hu6. For ^89^Zr labeling, MS-Hu6 was incubated with the chelator DFO-p-NCS for 3 hours, followed by incubation with ^89^Zr-oxalate for 1 hour at 37°C, ultrafiltration (cut off 10 kDa), and thin layer chromatography for quality check (29) (see Methods). ^89^Zr–MS-Hu6 was injected as a single dose of 250 µCi (∼250 µg) into the retroorbital sinus of 3–month–old male C57BL/6 mice (*N*=5 mice). For γ–counting few drops of blood were drawn from the tail vein at 5, 30 and 60 minutes, and then at 2, 4, 24, 48, 72 and 120 hours. There was an increase in serum ^89^Zr–MS-Hu6 levels to a C_max_ of 29.6 µg/mL, which was followed by a gradual decay of radioactivity with a β phase t_½_ of 32 hours (Fig. 4A).

**Figure 4:**
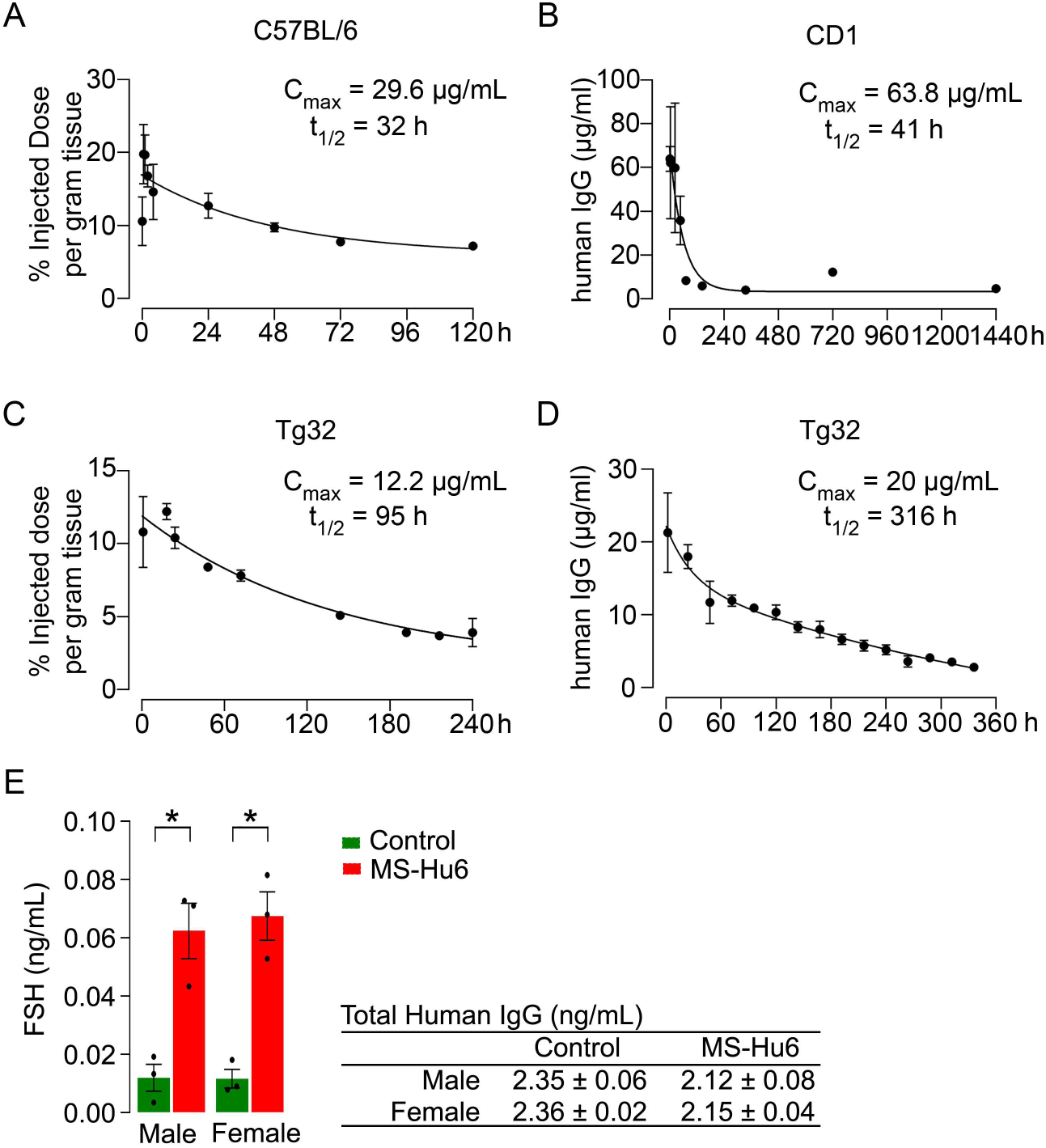
Pharmacokinetics and Target Engagement of MS-Hu6. Plasma levels, C_max_ and β phase t_½_ values for MS-Hu6 injected into C57BL/6 mice (250 µCi of ^89^Zr–MS-Hu6, *N*=5 mice followed longitudinally) **(A)**, CD1 mice (200 µg biotinylated MS-Hu6, *N*=3/time point) **(B)**, *Tg32* mice (250 µCi of ^89^Zr–MS-Hu6, *N*=5 mice followed longitudinally) **(C)** and *Tg32* mice (200 µg unconjugated MS-Hu6, *N*=3/time point) **(D)**. **(E)** Endogenous murine FSH in male and female mouse serum bound to injected MS-Hu6 *versus* that bound to control human IgG (hIgG). Detection of the MS-Hu6–FSHβα complex was achieved using an in–house ELISA in which the plate was coated with anti–human Fc and the complex captured by an antibody to the α subunit of FSH (*N*=3 biologic replicates). Of note is that total human IgG (control IgG or MS-Hu6) captured by an in–house ELISA was not different across treatment groups (E, Table).

As C57BL/6 mice are inbred strains, we attempted to validate the pharmacokinetic studies in an outbred strain—CD1. The latter mice display genetic diversity reminiscent of the human population, and are used widely for toxicology and efficacy testing (30). For biotinylation, MS-Hu6 was incubated in the presence of NHS ester–biotin in NaHCO_3_ (pH 8) and the product was purified through ultracentrifugation (cut off 10 kDa) (see Methods). Biotinylated MS-Hu6 (200 µg) was injected intraperitoneally with blood sampling by cardiac puncture at 2, 4, 24, 48, 72, 144, 336, 720, and 1440 hours (*N*=3 mice *per* time point), and ELISA–based measurements through capture by streptavidin-HRP. This yielded a C_max_ of 63.8 µg/mL and a β phase t_½_ of 41 hours (Fig. 4B).

To enable mouse–to–human comparisons, we studied the pharmacokinetics of MS-Hu6 in *Tg32* mice. These mice express the *FCGRT* transgene encoding the human FcRn receptor on chromosome 2 on a *Fcgrt^-/-^* background. *Tg32* mice show decreased plasma clearance of *human* IgG–based therapeutics––thus, more closely mimicking human pharmacokinetics than C57BL/6 mice. We injected 3–month–old male *Tg32* mice with ^89^Zr–MS-Hu6 (∼250 µCi or 250 µg) into the retroorbital sinus followed by sampling at 1, 18, 24, 48, 72, 144, 192, 216 and 240 hours (*N*=5 mice). Expectedly, the β phase t_½_ increased to 4 days (95 hours), with a C_max_ of 12.2 µg/mL (Fig. 4C). We further studied the profile of i.p. administered unconjugated MS-Hu6 in *Tg32* mice by injecting a single bolus dose of 200 µg, and measuring human IgG by an in–house sandwich ELISA in which anti–human Fc and Fab were used to capture and detect bound MS-Hu6, respectively (*N*=3 mice). This yielded C_max_ of 20 µg/mL, with β phase t_½_ of 13 days (316 hours) (Fig. 4D). Of note is that the β phase t_½_ for i.v. trastuzumab, currently in human use, is ∼8.5 days in *Tg32* mice (31)––this latter β decay translates into 21–day dosing intervals.

We looked for engagement of circulating MS-Hu6 with its target—FSH. For this, we injected groups of male and female C57BL/6 mice, intraperitoneally, with 200 µg MS-Hu6 or human IgG. Mice were bled 16 hours later and total IgG (mouse and human) was pulled down using protein A beads. The content of human IgG (control IgG or MS-Hu6) in the eluate, measured by a sandwich ELISA in which plates were coated with anti–human Fc and an anti– human IgG for capture, remained unchanged. We then subjected the same eluate to an sandwich ELISA, in which the plate was coated with anti–human Fc, and the MS-Hu6–FSHβα complex was captured by an antibody to the α subunit of FSH (incubation at 4°C, overnight). We found that injected MS-Hu6, but control human IgG bound endogenous FSH *in vivo* (Fig. 4E).

### Biodistribution and Excretion of MS-Hu6

To study the biodistribution of ^89^Zr–MS-Hu6, we performed PET-CT scanning of the above treated mice at 24, 48 and 72 hours (*N*=3 mice). The images show evidence of both decay of circulating radioactivity after 24 hours, and its persistence in multiple organs up to 72 hours (Fig. 5A). Maximal retention in terms of standardized uptake values (SUVs, normalized to muscle) was noted in the liver, with persistence in regions of interest, namely bone marrow, subcutaneous and visceral WAT depots, and the brain region (Fig. 5B). Of note is that these values reflect the presence of ^89^Zr–MS-Hu6 in both blood and tissues.

**Figure 5:**
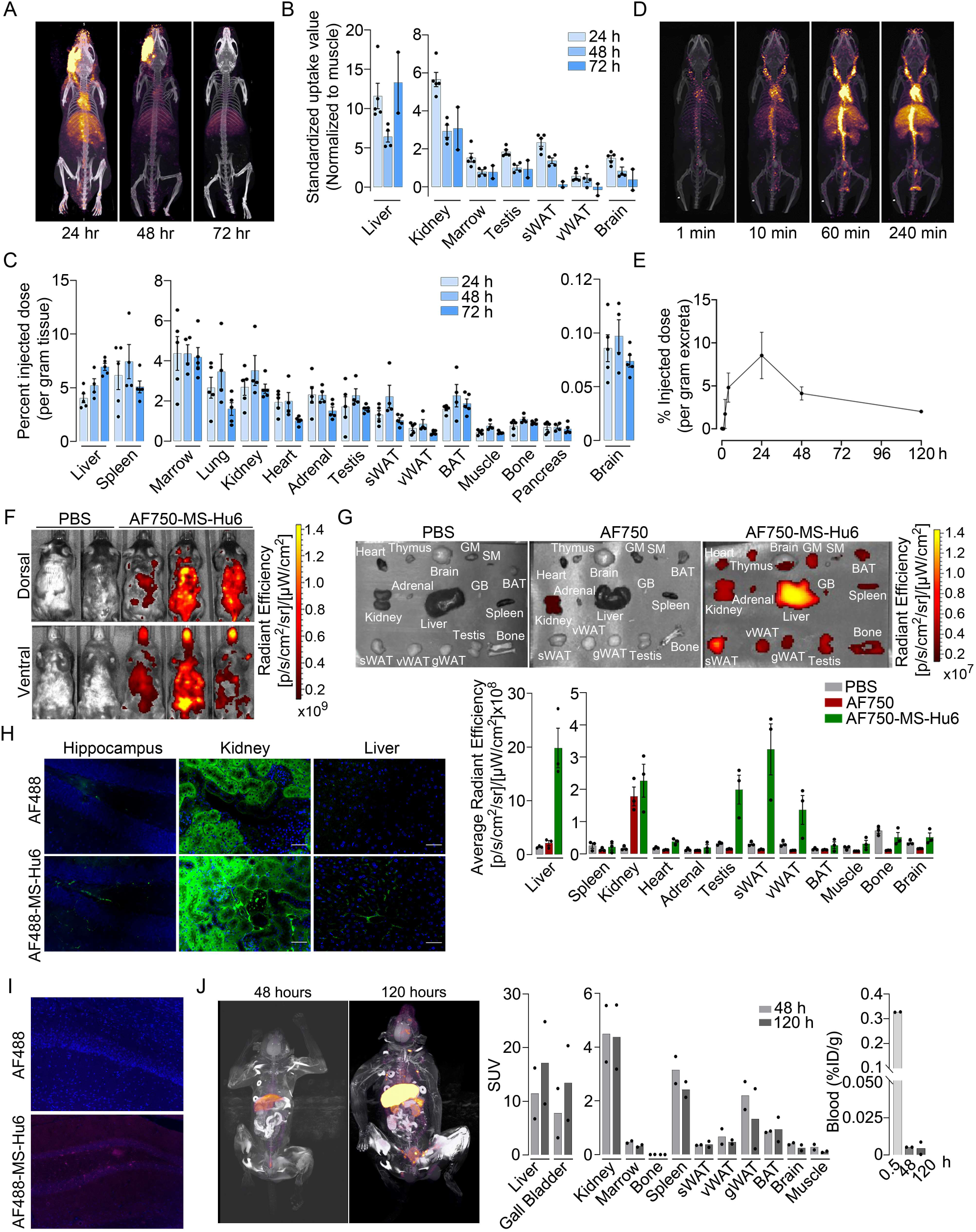
Biodistribution and Excretion of MS-Hu6 in Mice and Monkeys. Representative PET-CT images of mice treated with a single bolus dose of ^89^Zr–labeled MS-Hu6 (250 µCi) at 24, 48 and 72 hours **(A)**, together with quantitation in terms of standardized value uptake units (SUVs, normalized to muscle) in different organs (*N*=5, 4 and 2 mice for the three time points, respectively) **(B)**. ^89^Zr–MS-Hu6 (γ–counts) in individual tissues isolated following perfusion of the mice with 20 mL PBS (*N*=5, 4 and 5 mice for the three time point, respectively) **(C)**. Dynamic PET/CT images showing the uptake of ^89^Zr–MS-Hu6 over 240 minutes **(D)**. Time course of excretion of ^89^Zr–labelled MS–Hu6 in feces (*N*=5 mice/time point) **(E)**. Emitted whole body radiance on IVIS imaging of C57BL/6 mice injected with AF750-MS-Hu6 (200 µg) or PBS **(F)**. IVIS imaging and quantitation (average radiance) of isolated perfused tissues, as shown. following AF750-MS-Hu6, AF750 or PBS injection (*N*=3 mice/group) **(G)**. Immunofluorescence micrographs of hippocampal, kidney and liver sections from C57BL/6 mice injected with AF488-MS-Hu6 or unconjugated AF488 (200 µg/mouse, i.v.) **(H)**. Counterstaining with an anti–human IgG confirmed localization **(I)**. Whole body PET/CT image, quantitation (SUVs) of multiple organs and serum radioactivity (γ–counts) following a single i.v. injection of ^89^Zr-MS-Hu6 ((1.3 mg, ∼1.3 mCi)) in two *Cynomolgus* monkeys aged 14 and 15 years, respectively **(J)**.

To determine the extent to which of ^89^Zr–MS-Hu6 persisted in individual tissues, we perfused the mice with 20 mL PBS before sacrifice and tissue isolation for γ–counting. Significant concentrations of ^89^Zr–MS-Hu6 were detected in multiple organs, including bone, bone marrow, subcutaneous WAT, visceral WAT, and BAT (Fig. 5C). Minimal amounts of ^89^Zr–MS-Hu6 were detected in isolated brain tissue at 72 hours—this is consistent with the low penetration of IgGs into the brain (0.05 to 0.1%) (Fig. 5C). To hone into the early events, we monitored the uptake of ^89^Zr–MS-Hu6 by dynamic PET-CT imaging over 240 minutes. At 10 minutes, radioactivity was detected mainly in large vessels, which was followed at 60 and 240 minutes by permeation into organs (Fig. 5D). As would be expected, radioactivity was not detected in the urine, but instead appeared in the feces (Fig. 5E).

To complement the ^89^Zr–based biodistribution studies, we labelled MS-Hu6 with Alexa-Fluor-750 (AF750), and injected C57BL/6 mice (*N*=3 mice) intravenously through the tail vein with AF750–MS-Hu6 (200 µg), AF750 alone or PBS. At 16 hours post–injection, anaesthetized mice were imaged using the IVIS platform. We found significant soft tissue distribution of AF750–MS-Hu6 (Fig. 5F). The mice were then perfused with PBS, followed by IVIS imaging of isolated tissue. Consistent with the ^89^Zr-based studies, there was uptake of AF750–MS-Hu6 by liver, kidney, fat depots, bone, and brain (Fig. 5G). In contrast, in the AF750 (dye only) control group, localization was noted only in the kidney due to dye excretion, and not in other organs—in all, confirming organ retention of the AF750–MS-Hu6, and excretion of the unconjugated dye. Because of the expected minimal localization of MS-Hu6 in the brain, we further performed confirmatory studies by immunofluorescence. For this, we injected AF488–MS-Hu6 or unconjugated AF488 into the tail veins of C57BL/6 mice (200 µg *per* mouse). We detected immunofluorescence in liver, kidney and hippocampal sections in mice treated with AF488–MS-Hu6 (Fig. 5H). In contrast, AF488– treated mice showed fluorescence in kidney sections, but not in liver or brain (Fig. 5H). Staining of hippocampal sections with an anti–human IgG confirmed localization (Fig. 5I).

To understand MS-Hu6 biodistribution as it may apply to humans, we injected ^89^Zr-MS-Hu6 as a single bolus dose (1.3 mg, ∼1.3 mCi) into the tail veins of two male *Cynomolgus* monkeys aged 14 and 15 years, respectively. Blood was drawn *via* tail vein at 5 minutes and at 48 and 120 hours. ^89^Zr-MS-Hu6 peaked in the blood at 5 minutes, with an expected decline, *albeit* with persistence in the serum, at 48 and 120 hours (Fig. 5J). PET/CT scanning revealed high SUV values in the liver and gall bladder, with lower SUVs in the kidney, spleen, fat depots, bone marrow, and the brain area (Fig. 5K).

### Acute and Chronic Safety of FSH Blockade

We monitored standard safety parameters in treated monkeys up to 100 minutes, and did not observe significant acute or delayed changes post–injection in heart rate, respiratory rate, mean arterial blood pressure, systolic or diastolic blood pressure, or rectal temperature (Fig. 6A). We also drew bloods at day 0 (pre–injection) and at days 2 and 5 post–injection. No concerning deviations from normative values were noted (Fig. 6B). This suggests that, *albeit* at a low dose, MS-Hu6 as a single intravenous bolus injection into monkeys appeared to be generally safe.

**Figure 6:**
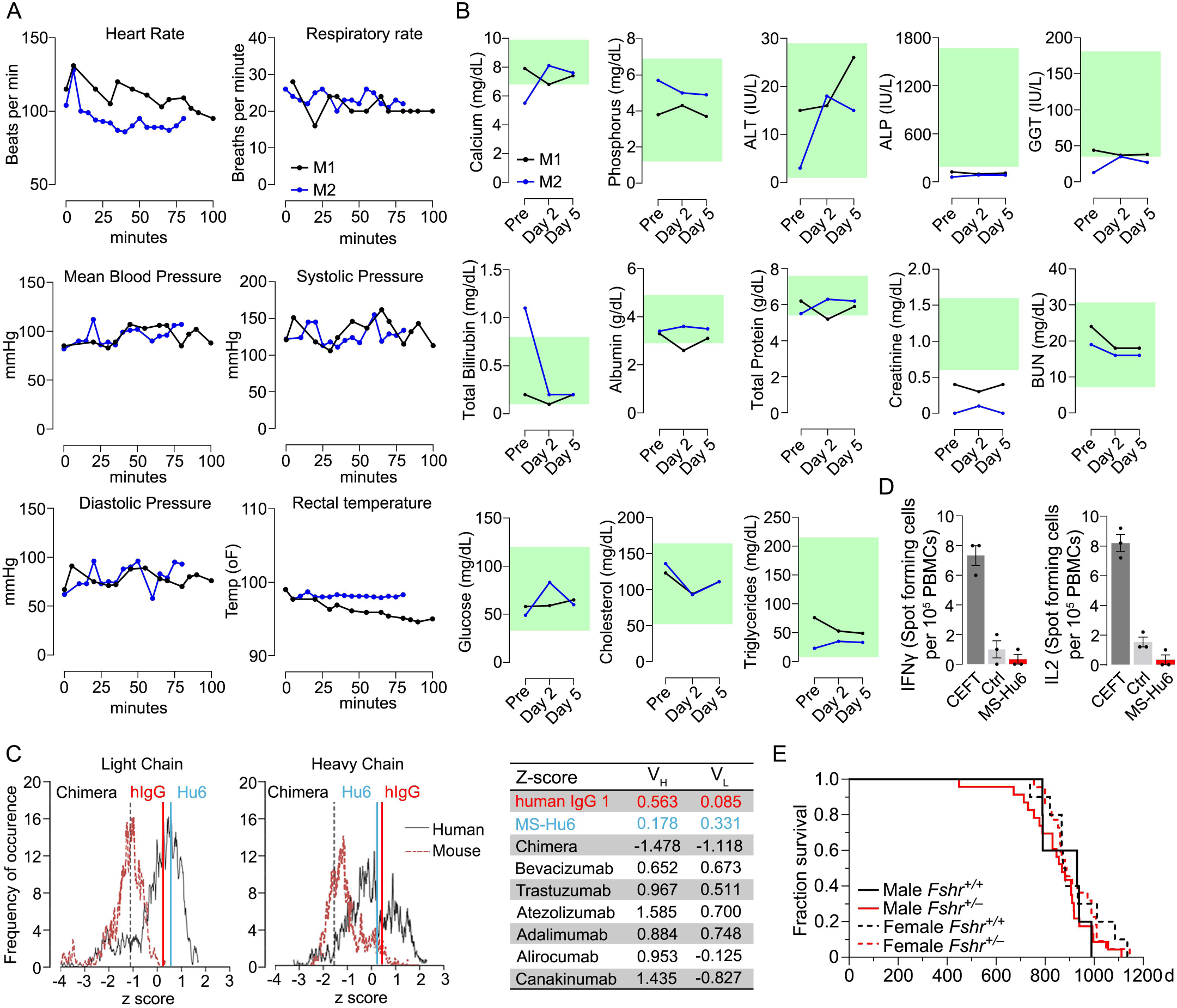
Acute and Chronic Safety of MS-Hu6. Effects on physiological parameters (monitored up to 100 minutes **(A)** and serum biochemistry (at days 0, 2 and 5) **(B)** after injecting ^89^Zr-MS-Hu6 as a single i.v. bolus dose (1.3 mg, ∼1.3 mCi) into the tail veins of *Cynomolgus* monkeys (Andy, 14 years/9.8 kg, and Scott, 15 years/6.15 kg). Normative serum biochemistry data (green) from: (65). *In silico* assessment of “humanness” using abYsis, with *Z*-scores comparing humanized MS-Hu6, mouse–human chimeric antibody (26), and human IgG1 **(C)**. Table shows narrowly distributed *Z*-scores after inputting primary sequences of FDA–approved, clinically–utilized humanized (“zumab”) and fully human (“mab”) antibodies: bevacizumab (anti– VEGF, Avastin®); trastuzumab (anti–Her2, Herceptin®); atezolizumab (anti–PD-L1, Tecentriq®); adalimumab (anti–TNFα, Humira®); alirocumab (anti PCSK9, Praluent®); and canakinumab (anti–IL-β; Ilaris®) (C). Assessment of immunogenicity using ELISPOT assays for the inflammatory cytokines IL-2 and IFNγ in human peripheral blood mononuclear cell cultures in response to added MS-Hu6, DMEM (Ctrl), or CEFT peptide pool (positive control, Immunospot) **(D)**. Kaplan–Meier survival curves showing that haploinsufficiency of *Fshr*, which otherwise mimics the effect of MS-Hu6 on body fat and bone mass (*c.f.* Fig. 1C) (13, 16) does not reduce lifespan compared with wild type littermates over 1100 days (3 years) (5, 10 mice for wild type male and female, and 23, 22 mice for male and female *Fshr*^+/-^ mice, respectively) **(E)**.

MS-Hu6 was generated by swapping the mouse framework region of our parent mouse monoclonal antibody Hf2 with the human IgG1 framework, keeping the CDR itself unaltered. While both Fc and Fab region are human, mutations were introduced in the framework flanking the CDR region. Because MS-Hu6 is not fully “human”, we first determined its “humanness” *in silico* by inputting the V_L_ and V_H_ sequences into abYsis. Comparison of *Z*-scores revealed a right– shift in comparison with the human–mouse chimeric antibody (26), and in correspondence with human IgG1 (Fig. 6C). In addition, we inputted the primary amino acid sequences of commercially utilized humanized (“zumab”) and fully human (“mab”) to find that the *Z*-scores fell in a narrow range away from our chimera or mouse IgGs (Fig. 6C, Table). We next tested immunogenicity experimentally using ELISPOT. The production of inflammatory cytokines IL-2 and IFNγ in human peripheral blood mononuclear cell cultures was unaltered by MS-Hu6, in comparison with a standard CEFT peptide pool (positive control, Immunospot) (Fig. 6D). Finally, we provide genetic evidence, using our *Fshr*^+/-^ mouse, that haploinsufficiency of the FSHR—which mimics the effect of FSH blockade on obesity and osteoporosis (13, 15, 16)—does not affect lifespan negatively in male or female mice (Fig. 6E).

### Developability, Formulation and Physicochemistry of MS-Hu6

Therapeutic antibodies selected on the basis of affinity, potency, specificity, functionality and pharmacokinetics might not have unsuitable physicochemical attributes making it difficult to streamline optimal manufacturing. It is therefore imperative that, at an early stage, we determine physicochemical properties, after a rough *in silico* check for ‘red flags’ (32–34). We thus used a computational tool, Protein–Sol, based on machine learning of amino acid sequences and physicochemical variables from 48 FDA–approved antibodies and 89 antibodies in late–stage clinical development (https://protein-sol.manchester.ac.uk/abpred) (32–34). In the initial iteration, Protein–Sol provided predicted values for 12 separate physicochemical parameters that determine manufacturability. For all outputs, after inputting V_H_ and V_L_ regions, MS-Hu6 fell within acceptable thresholds, and was therefore deemed to be “safe” (Fig. 7). In essence, the physicochemical properties of MS-Hu6 were likely to be broadly similar to FDA–approved antibodies.

**Figure 7:**
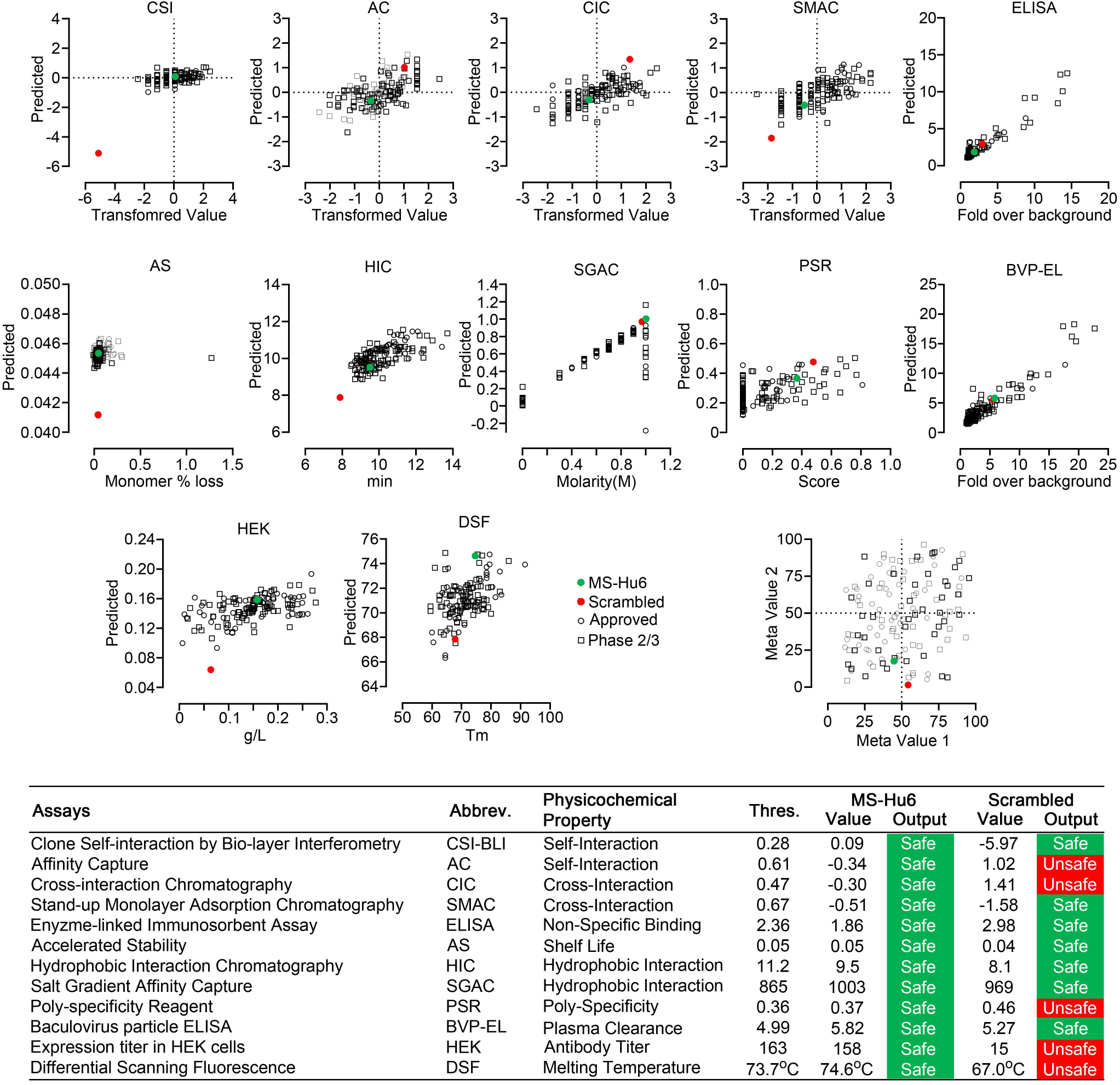
Manufacturability of MS-Hu6. **(A)** Protein–Sol was used to compare twelve physicochemical parameters that were computationally–derived for MS-Hu6 against experimental derivations of 48 FDA–approved antibodies and 89 antibodies in late–stage (phase 2/3) clinical development. MS-Hu6 fell within the “safe” range and was therefore considered manufacturable. A variation of MS-Hu6, in which the CDR region was scrambled, failed 5 of 12 outputs, indicating non–manufacturability. While 65% FDA–approved monoclonal antibodies show no ‘red flags’, those with up to 4 red flags have been approved by the FDA (34). Also shown are meta values for both MS-Hu6 and its scrambled sequence (1=best; 100=worst) derived by averaging ranks for 8 experimental parameters. MS-Hu6 fell within the lower left quadrant, confirming that physicochemical properties were acceptable for manufacturing.

For validation, we inputted a version of MS-Hu6, wherein the CDR region was scrambled—5 of 12 outputs, namely affinity capture, cross interaction chromatography (CIC), polyspecificity reagent (PSR), expression titers in HEK cells (HEK), and differential scanning fluorescence (DSF), fell outside the respective thresholds—an early indication that the scrambled version was not manufacturable (Fig. 7). In fact, while 65% FDA–approved monoclonal antibodies show no ‘red flags’, those with up to 4 red flags have been FDA–approved (34). To complement data from individual outputs, we derived a meta value for both MS-Hu6 and its scrambled sequence (1=best; 100=worst) by averaging ranks for 8 experimental parameters. We found that meta value pairs fell within the lower left quadrant, suggesting overall acceptable physicochemical properties, even in comparison with certain FDA–approved antibodies in the upper right quadrant (Fig. 7).

Before testing the physicochemical characteristics of MS-Hu6 experimentally, we created an optimal formulation. To prevent deamidation and isomerization at neutral and basic pHs, therapeutic antibodies are generally formulated at pHs away from their isoelectric pH (pI) (35, 36). Using Expasy, we predicted the pI for MS-Hu6 as 8.58. Isoelectric focusing confirmed a pI pf 8.7 (Fig. 8A). We tested 215 combinations of salt, detergent and sugars for their thermal stability (not shown). This yielded in a near–final formulation for MS-Hu6—stock solution of 2 mg/mL in 20 mM phosphate, 0.001% (v/v) Tween-20, 1 mM NaCl, and 260 mM sucrose (pH = 6.58). We found that both Fc and Fab regions of the formulated MS-Hu6 showed a thermal shift (ΔT_m_) compared with MS-Hu6 in PBS—and, hence, was confirmed as being more stable (Fig. 8B). We also examined the binding of formulated MS-Hu6 to purified human FSH. A ΔT_m_ of 3.1 °C of the Fab, and not the Fc region, established greater thermal stability due to FSH binding (Fig. 8C). Furthermore, and of note, is that compared with MS-Hu6 in PBS, formulated MS-Hu6 showed dampened peak signals (Fig. 8B). The latter finding indicates that a lower number of antibody molecules underwent unfolding—implying enhanced stability—despite identical added concentrations (20 µg/well).

**Figure 8:**
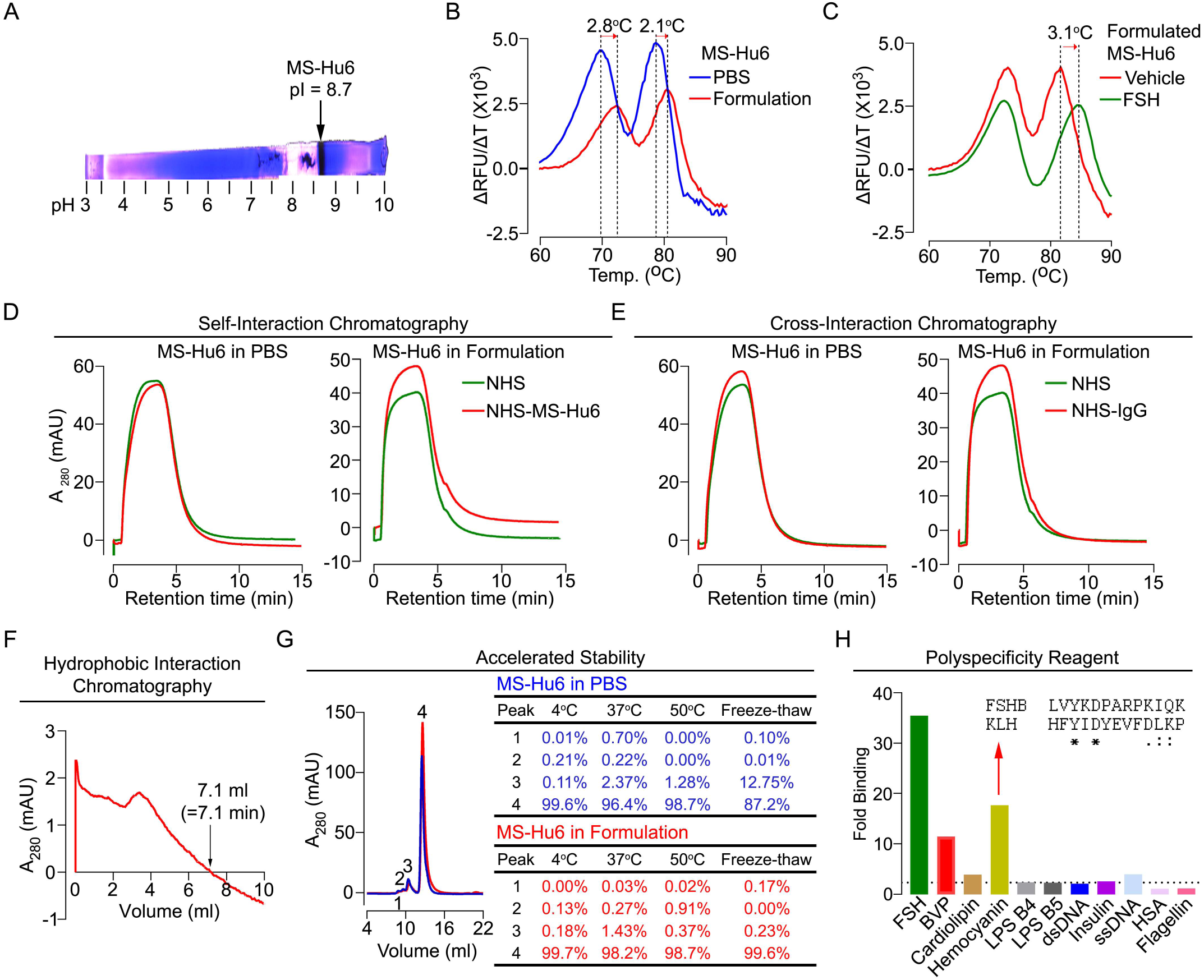
Physicochemical Characteristics of MS-Hu6. Isoelectric focusing confirmed a pI pf 8.7 for MS-Hu6, consistent with its *in silico* prediction (Expasy) of 8.58 **(A)**. Thermal shift assays were used to examine stability of both Fc and Fab regions of formulated MS-Hu6 *versus* MS-Hu6 in PBS **(B)**, as well as stability of FSH binding to the Fab region of formulated MS-Hu6 **(C)**. UV absorbance (280 nm) readout of self–interaction chromatography (SIC) **(D)** or cross–interaction chromatography (CIC) **(E)** to assess binding of formulated MS-Hu6 *versus* MS-Hu6 in PBS with self or human IgG, respectively. Hydrophobic chromatography showing UV absorbance (280 nm) of the eluate from a butyl sepharose column upon passing MS-Hu6 at pH 6.5 over 20 minutes at a flow rate of 1 mL/min (retention time shown) **(F)**. Representative size exclusion chromatograms and area under the peak for MS-Hu6 in PBS or formulated MS-Hu6 following stress testing by 3 cycles of freeze–thaw or incubation at 4 °C, 37 °C and 50 °C for 1 week **(G)**. Reactivity of MS-Hu6 to a standard panel of antigens, including cardiolipin, hemocyanin (KLH), lipopolysaccharides B4 and B5 (LPSB4 and LPSB5), double and single stranded DNA (dsDNA and ssDNA), insulin, human albumin, flagellin, and baculovirus particles (BVP) (ELISAs) **(H)**. Homology of KLH with the epitope against which MS-Hu6 was raised, showing identical (*) and conserved (:) amino acid residues (H).

For a full physicochemical characterization of the formulated MS-Hu6, we used a battery of biochemical tests, namely CIC, self–interaction chromatography (SIC), size exclusion chromatography (SEC), hydrophobic chromatography, and polyspecificity assays. For CIC and SIC, we created an in–house NHS ester column conjugated either with human IgG (for CIC) or MS-Hu6 (for SIC). MS-Hu6 was passed through an unconjugated column both in PBS and as a formulation, as well as through the two conjugated columns. Figs. 8D and 8E show that retention times of MS-Hu6 in the respected conjugated columns were not different from those of the unconjugated column—this confirms that MS-Hu6 does not appreciably interact with itself or with human IgGs, also indicative of little or no aggregation.

Highly soluble and hydrophilic monoclonal antibodies are expected to behave robustly during the manufacturing process. On the other hand, hydrophobic antibodies with high sensitivity to salt may display problems, such as poor expression, aggregation or precipitation during purification. Delayed retention times in a chromatographic assay with a hydrophobic matrix are indicative of a tendency for precipitation. We used a butyl sepharose column, and passed MS-Hu6 in 1.8 M NH_2_SO_4_ and 0.1 M Na_2_PO_4_ at pH 6.5 over 20 minutes at a flow rate of 1 mL/min. UV absorbance was monitored at 280 nm to yield a retention time of 7.1 minutes (Fig. 8F), which is below the theoretical threshold of 11.2 min (Fig. 7).

We stressed formulated or PBS–containing MS-Hu6 through three freeze–thaw cycles (from –80°C to room temperature), and by incubation for 1 week at 4 °C, 37 °C or 50 °C, followed, in all cases, by SEC (Fig. 8G). We noted a major peak (#4) and three minor high molecular weight peaks (#1 to 3). Areas under peaks 1 and 2 were between 0 and 0.91% of the total eluate under all conditions for both PBS–containing and formulated MS-Hu6. Peak 3 remained generally low (<1.5%) with formulated MS-Hu6, but was considerably higher (up to 12.7%) during freeze–thaw in PBS. Furthermore, the major peak 4 was consistently >99% with formulated MS-Hu6, particularly when compared to PBS–containing MS-Hu6 under freeze thaw (87.2%). In all, the formulation protected against aggregation even under extreme stress conditions. Extending the protocol to 25 mL elution yielded no fragment peaks under any condition (Fig. 8G).

Therapeutic antibodies must undergo a test for polyreactivity to a panel of relevant antigens, including cardiolipin, hemocyanin (KLH), lipopolysaccharide (LPS), double and single stranded DNA, insulin, human albumin, flagellin, and baculovirus particles (BVP). The fold change for MS-Hu6 binding to every molecule in ELISAs, except hemocyanin and BVP, was below a threshold of 3, and was therefore acceptable (Fig. 8H). BVP scores greater than 5 have been shown to enhance antibody clearance, with potential effects on t_½_. In contrast. binding to hemocyanin, an arthropod protein, often used as a carrier for synthetic peptides during immunization (37), was likely due its homology with the 13–amino acid sequence of FSHβ that binds MS-Hu6 (Fig, 8H). However, as hemocyanin was not used during the production of the parent monoclonal antibody, Hf2, this is an irrelevant finding.

## DISCUSSION

Not many biologics, or indeed new therapeutics, undergo robust preclinical evaluations in academic medical centers outside of the pharmaceutical or biotechnology space. Our comprehensive analysis of the biology and physicochemistry of a first–in–class, humanized FSH– blocking antibody, MS-Hu6, establishing it as efficacious, safe and manufactureable may arguably be amongst the first attempts at an extensive in–house effort. In 2003, we conceived the idea that pituitary hormones, including FSH, have direct actions on bone (13, 38–40); shifted this new paradigm to establish novel actions of FSH on fat, and more recently, on brain (16, 41); created the first polyclonal FSH–blocking antibody targeted to a 13–amino–acid–long receptor–binding epitope of FSHβ (17, 18); and finally developed a humanized multipurpose therapeutic (26) for future use in osteoporosis and obesity, and perhaps even in hyperlipidemia and Alzheimer’s disease. Notably, the in–house studies described here have been carried out using our own platform that complies with Good Laboratory Practice, as mandated by the Food and Drug Administration’s Code of Federal Regulations [Title 21, Chapter 1A, Part 58]. This means, that this Investigational New Drug (IND)–enabling dataset forms the basis of late stage development and future first–in–human studies.

It is now well recognized that targeting the receptor–binding epitope of FSHβ to block its interaction with the FSHR inhibits bone resorption, increases bone formation and bone mass, reduces body fat, and enhances thermogenesis (15–19, 22). Notably, these effects are triggered not only with our polyclonal and monoclonal FSH–blocking antibodies (15–18), but also through the use by others of vaccines, such as a GST–FSHβ fusion protein (19) or tandem repeats of the epitope (22). It is also important to note that even with high antibody doses, such as 200 µg/mouse/day, serum estrogen levels are unchanged (16, 17), likely because of the abundance of FSHRs in the ovary that remain responsive to lowered levels of bioavailable FSH. These preclinical data in mice establishing FSH as an actionable target are reinforced by striking estrogen–independent correlations in women between serum FSH, rapid bone loss and visceral obesity during the menopausal transition, at which time serum estrogen is relatively normal and FSH levels are rising (4, 5, 9–12). This window is likely the most opportune to prevent both bone loss and obesity through selective FSH blockade. In this regard, while negative data with GnRH modulators are mostly confounded by concomitant changes in LH and GnRH (14), it is clear that low gonadotropin levels in triptorelin–treated men with prostate cancer are associated with lower fat mass and body weight than men undergoing orchiectomy, wherein gonadotropins are high (23). Overall, therefore, the data together lend credibility to the idea that FSH–driven changes, at the very least in body composition, in both sexes can be rescued by blocking FSH. The selective inhibition of FSH action therefore becomes a worthy imperative.

Menopause is also associated with dyslipidemia, which has long been thought to result from estrogen deficiency. However, there is compelling epidemiologic evidence that high serum FSH levels correlate with serum total and LDL cholesterol in post–menopausal women, and importantly, that total cholesterol rises across the perimenopausal transition, essentially tracking closely with bone loss and obesity (5, 7, 25, 42). Impressively, exogenous FSH, in the presence of estrogen clamped at normal levels, increases serum total cholesterol in mice fed on a high cholesterol diet (24). And, consistent with the idea that FSH is an estrogen–independent driver of menopausal hypercholesteremia, an FSH–blocking antibody, which we have previously shown to be active in bone cells, lowers serum cholesterol (18, 24). There is also human evidence that reducing serum FSH by >30% from its zenith in post–menopausal women through estrogen replacement therapy lowers serum cholesterol (25). As in the case of bone and fat cells (13, 16), hepatocyte FSHRs couple with Gi_2α_, which signals through Akt to inhibit FoxO1 binding with the *Srebp2* promoter and prevent its repression. Upregulated *Srebp2*, which drives *de novo* cholesterol biosynthesis, results in increased cholesterol accumulation and release (24). This action is in addition to the lowering of LDLR expression by FSH (25). Notwithstanding cholesterol–lowering mechanism(s), which are likely to be explored even further, it is possible that an FSH–blocking therapeutic, such as MS-Hu6, could have additional actions on lipid metabolism in people.

Furthermore, what underpins the preponderance of Alzheimer’s disease (AD) in post– menopausal women, particularly in relation to disease risk, progression and severity, has remained unclear, now for decades. A role for post–menopausal hypoestrogenemia remains controversial, with improvement (43), no change (44, 45) or worsening (46) of cognition with estrogen replacement therapy. In contrast, high serum FSH is strongly associated with the onset of AD, and has thus been suggested as a possible mediator (47, 48). More importantly, certain neuropathologic features, including neuritic plaques, neurofibrillary tangles, and gliosis often begin during the perimenopausal transition (49–53). During this period, women also show a sharp decline in memory function and increased risk of mild cognitive impairment and dementia (11, 52, 53). We have recently documented exaggerated AD pathology and cognitive decline upon ovariectomy or exogenous FSH injection in three murine models of AD, even in the face of estrogen levels clamped in the normal range (41). This phenotype arises from the action of FSH on hippocampal and cortical neuronal receptors using a pathway involving CEBP/β and the δ– secretase asparagine endopeptidase (41). Most notably, however, we found that our polyclonal FSH–blocking antibody, which shares the target epitope with MS-Hu6 (17, 26), prevented the AD–like phenotype induced upon ovariectomy (41)—providing a clear avenue for further exploration of the effects of MS–Hu6 in models of AD.

In all, therefore, we and others have unraveled new actions of FSH that assign it as an actionable target requiring a highly specific approach to block its action in people. We believe that MS–Hu6 with an affinity approaching that of trastuzumab (54), which we have carefully characterized in terms of structure, biological actions, pharmacokinetics, target engagement, biodistribution, safety and manufacturability, is poised for future testing in human trials. Admittedly ambitious, we envisage, in a best case scenario, and if mouse data translates into people, of treating four diseases that affect millions of women and men worldwide—namely obesity, osteoporosis, dyslipidemia and neurodegeneration—with a single multipurpose FSH– blocking agent.

## ACKNOWLEDGEMENTS

Work at Icahn School of Medicine at Mount Sinai performed at the Center for Translational Medicine and Pharmacology was supported by R01 AG071870 to M.Z., T.Y. and S.-M.K.; R01 AG074092 and U01 AG073148 to T.Y and M.Z.; U19 AG060917 to M.Z. and C.J.R.; and R01 DK113627 to M.Z. and J.I. Work at U.S. Department of Agriculture, Agricultural Research Service, Grand Forks Human Nutrition Research Center (USDA ARS GFHNRC) was supported by the Project Plan #3062-51000-053-00D to J.J.C. M.Z. also thanks the Harrington Discovery Institute for the Innovator–Scholar Award towards development of the FSH Ab. C.J.R. acknowledges support from the NIH (P20 GM121301 to C.J.R). Mention of trade names or commercial products in this publication is solely for the purpose of providing specific information and does not imply recommendation or endorsement by the U.S. Department of Agriculture. USDA is an equal opportunity provider and employer. The findings and conclusions in this manuscript are those of the authors and should not be construed to represent any official USDA of U.S. Government determination or policy.

## AUTHOR CONTRIBUTIONS

M.Z., and T.Y. designed the experiments; S.G. performed and/or oversaw most experiments with T.-C.K. and F.K.; D.S., C.R., T.-C.K. and S.G created the and tested the antibody formulation; A.G., S.M. and V.R. carried out histology and histomorphometry; V.D. carried out the bone treatment protocol under C.J.R.’s supervision (Maine Medical Center research Institute, Scarborough, ME); G.P., J.M., J.C.F., T.G.J.M.P., A.T., M.v.L., and Z.F. together with S.G. and T.-C.K. designed and executed experiments involving ^89^Zr labeling, PET/CT scanning and analyses, and biodistribution and safety studies in monkeys, and J.M. performed PET/CT and PET/MRI data analyses; J.N., E.S., S.G. H.K., K.I. and M.B. assisted with the mouse studies on body composition under the guidance of S.G.; T.-C.K., P.K., A.P., and S.G.. performed biodistribution studies in mice; T.-C.K., F.K. and S.G. carried out pharmacokinetic analyses; J.C. performed the µCT data analysis (USDA ARS GFHNRC, Grand Forks, ND) F.S., K.S. and J.N. performed MRI studies; R.B. and V.M. performed studies using *Fshr*^-/-^ mice; S.G. and T.-C.K. performed *in silico* developability analyses; S.G., D.S. and T.-C.K. performed physicochemical analyses; S.G., N.Z. and T.Y. performed and interpreted *in silico* immunogenicity and cell proliferation and viability assays; M.S. and M.M. provided regulatory expertise on constructing the GLP platform; L.C. and S.B. drafted protocols using GLP standards; A.M., V.R. and J.I. maintained and provided quality control for the GLP platform; D.L. and S.-M.K. assisted with checking datasets independently and assuring data provenance; J.C. and M.M. provided advice on IND enablement in relation to preclinical studies; M.I.N., C.J.R, T.Y. and M.Z. composed and edited the manuscript.

## METHODS

### Animals

Colonies of male and female C57BL6 mice, and male *Tg32*, *Fshr^+/-^* mice, and ThermoMice were obtained from Jackson Labs. Male CD1 mice were from Charles River Laboratories. Mice were maintained in–house at Icahn School of Medicine at Mount Sinai and/or Maine Medical Center Research Institute. They were either fed on normal chow or on a high–fat diet (DIO Formula D12492, 60% fat; Research Diets, Inc., New Brunswick, NJ.), with access to water *ad libitum*. The mice were housed in climate–controlled conditions with standard 12–hour light/dark cycles (6 AM to 6 PM). Nonhuman primates, namely *Cynomolgus* monkeys (*Macaca fascicularis*), were fed Teklad Global 20% Protein Primate Diet. All protocols were approved by Institutional Animal Care and Use Committees of Icahn School of Medicine at Mount Sinai and Maine Medical Center Research Institute.

### Body Composition

To study the effects of MS-Hu6 on body composition, we used male SV129 ThermoMice that were allowed *ad libitum* access to a high–fat diet and injected with MS-Hu6 or human IgG (7 µg/day, 5 days–a–week, i.p.) for 8 weeks. The latter dose was based on the *in vitro* IC_50_ of MS- Hu6, which was ∼30–fold lower than our polyclonal Ab (26). We determined net food intake and measured body weight weekly, and performed quantitative nuclear magnetic resonance (qNMR) at 8 weeks. For this, live mice were placed into a thin–walled plastic cylinder, with freedom to turn around. An Echo3-in-1 NMR analyzer (Echo Medical) was used to measure fat, lean and total mass, *per* manufacturer.

In ThermoMice, a luciferase reporter construct, *Luc2-T2A-tdTomato*, is inserted into the *Ucp1* locus on the Y–chromosome (55). Activation of *Ucp1* expression leads to upregulation of *Luc2*, which can be quantitated *in vivo* by radiance measurements using IVIS Spectrum *In Vivo* Imaging System (Perkin Elmer) following the injection of D-Luciferin (150 mg/kg). At 4 and 8 weeks, the mice were injected with D-Luciferin and radiance was captured from dorsal and ventral surfaces for optimal visualization of interscapular (BAT–rich) and inguinal (WAT–rich plus testes) regions, respectively. Isolated fat depots were weighed manually, and RNA extracted for quantitative PCR (qPCR) for fat (*Pparg*, *Fabp4* and *Cebpa*) and beiging (*Ucp1*, *Cox 8a*, *Cox7*, *Cidea*, and *Prdm16*) genes using appropriate primer sets and Prism 7900-HT (Applied Biosystems Inc.) (56).

### Histology and Immunodetection Methods

Tissues were subject to hematoxylin/eosin staining, immunohistochemistry for UCP1 (primary anti–UCP1 antibody: Abcam, Catalog# ab209483; secondary antibody: goat anti–rabbit IgG, Invitrogen, Catalog# 32260), or fluorescence for AF488 (conjugated to MS-Hu6) or AF750 (conjugated to human IgG1) [human IgG, Sigma Cataog # I12511; Alexa Fluor 488 Antibody Labelling Kit, Invitrogen, Catalog # A20181; SAIVI Rapid Antibody Labelling Kit, Alexa Fluor 750, Invitrogen, Catalog # S30046]. Frozen brain sections or formalin–fixed, paraffin–embedded sections were examined using an Observer-Z1 microscope (Zeiss, Germany We used the following ELISA kits: FSH and LH (MILLIPLEX MAP Mouse Pituitary Magnetic Beads, Catalog #: MPTMAG-49K, CLHMAG and RFSHMAG, respectively), testosterone (R and D Systems, Catalog #: KGE010), and activin A (ThermoFisher, Catalog # EM3RB).

### Bone Phenotyping

At Maine Medical Center Research Institute, BMD was measured by dual energy X-ray absorptiometry (DXA) (*Piximus*, Lunar) with a precision of <1.5% (57). At Mount Sinai, we used Osteosys iNSIGHT and analyzed the DXA data using Insight v1.0.6. In both instances, anaesthetized mice were subject to measurements, with the cranium excluded. The instruments were calibrated each time before use *per* manufacturer’s recommendation.

For µ–CT measurements, femoral epiphyses were scanned non–destructively by using a Scanco µCT scanner (µCT-40; Scanco Medical AG, Bassersdorf, Switzerland) at 12 µm isotropic voxel size, with X-ray source power of 55 kV and 145 µA, and integration time of 300 milliseconds. Trabecular microstructure was evaluated after removing the noise from the scanned grey–scale images using a low-pass Gaussian filter. A fixed bone mineral density threshold of 220 mg/cm^3^ was used to extract the mineralized bone from soft tissue and the marrow phase. Reconstruction and 3D quantitative analyses were performed using software provided by Scanco. The same settings for scan and analysis were used for all samples. Trabecular bone parameters included fractional bone volume (BV/TV), trabecular thickness (Tb.Th), trabecular number (Tb.N), trabecular spacing (Tb.Sp), and connectivity density (Conn.D).

Two–dimensional histomorphometry was performed in femoral epiphysis. Frozen non– decalcified sections (6–8 µm) were stained with a von Kossa staining kit (American MasterTech, Catalog # KTVKO), *per* manufacturer’s procedure. This provided measures of fractional bone volume (B.Ar/T.Ar), Tb.N, Tb.Sp and Tb.Th. Bone formation was quantified by dynamic histomorphometry following sequential injections of calcein (15 mg/kg) followed by xylelol orange (90 mg/kg) eight days apart, with the last injection 4 days prior to sacrifice. Parameters included mineral apposition rate (MAR) and bone formation rate (BFR),. Osteoclast resorbed surface (Oc.S/BS) was measured following TRAP staining with Leukocyte Acid Phosphatase TRAP Kit (Sigma Aldrich, catalog # 387A-1KT). RNA was extracted from isolated whole femur for qPCR for osteoblast (*Col1a1*, *Alpl*, *Runx2* and *Bglap*) and osteoclast (*Acp5*) genes using appropriate primer sets using Prism 7900-HT (Applied Biosystems Inc.) (56).

### Pharmacokinetics and Biodistribution in Mice

We prepared ^89^Zr–MS-Hu6 by incubating MS-Hu6 first with the chelator DFO-p-NCS for 3 hours at 37 °C (in steps of 5 µL until a 10–fold molar excess of chelator was achieved). The DFO-functionalized MS-Hu6 was washed 3 times with PBS in a 10 kDa ultracentrifugation tube before radiolabeling. ^89^Zr-oxalate was diluted with PBS and neutralized with 1M Na_2_SO_4_ before adding to functionalized MS-Hu6. This was followed by incubation with ^89^Zr-oxalate for 1 hour at 37 °C, ultrafiltration (cut off 10 kDa), and thin layer chromatography (with 50 mM EDTA) for quality check (29). C57BL/6 or *Tg32* mice were injected in separate experiments with ^89^Zr–MS-Hu6 as a single dose of ∼250 µCi (∼250 µg, 250 ± 40 µCi) into the retroorbital sinus. Timed blood (few drops drawn from the tail vein) and excreta collection was followed by weighing and γ–counting (PerkinElmer Wizard 2480 Automatic Gamma Counter, PerkinElmer, Waltham, MA). Values were corrected for decay and expressed as a percentage of injected dose *per* gram of tissue (%ID/gram).

In complementary experiments, MS-Hu6 was biotinylated by incubating with NHS ester– biotin (100 µg *per* mg MS-Hu6, dissolved in DMSO, Sigma, Catalog # H1759) for 4 hours at room temperature. NaHCO_3_ (1 M) was then added for 10 minutes at room temperature (pH 8) and the product was purified through ultracentrifugation (cut off 10 kDa). Biotinylated MS-Hu6 (200 µg) was injected i.p. into CD1 mice that underwent cardiac puncture for blood sampling and sacrifice at timed intervals. We used an in–house ELISA in which the plate was coated with individual sera and biotinylated MS-Hu6 was captured by streptavidin-HRP (Millipore, Catalog # 18-152). In a final experiment, unconjugated MS-Hu6 was injected i.p. into *Tg32* mice (200 µg). Serum was collected from groups of mice and an in–house assay was used in which anti–human IgG Fc (Sigma, Catalog # FAB3700259) was used to capture human IgG (MS-Hu6), with detection of the complex with goat anti–human IgG Fab (200 µg/well; Invitrogen, #31122).

For imaging, ^89^Zr–MS-Hu6–treated mice (above) were anesthetized using 1% isoflurane in O_2_ at a flow rate of ∼1.0 L/minute. PET/CT scans were performed using a Mediso nanoScan PET/CT (Mediso, Budapest, Hungary). For whole body CT scans, we used the following parameters: energy, 50 kVp; current, 180 µAs; and isotropic voxel size, 0.25 mm)—this was followed by a 30–min PET scan. Image reconstruction was performed with attenuation correction using the TeraTomo 3D reconstruction algorithm from the Mediso Nucline software (version 3.04). The coincidences were filtered with an energy window between 400 and 600 keV. Voxel size was isotropic with 0.4–mm width, and the reconstruction was applied for four full iterations, six subsets *per* iteration. Image analysis was performed using Osirix MD, version 11.0. Namely, whole body CT images were fused with PET images and analyzed in an axial plane. Regions of interest (ROIs) were drawn on various tissues. Testis, visceral WAT, subcutaneous WAT, kidneys, liver, and brain were traced in their entirety, and bone marrow uptake was assessed using three vertebrae in the lumbar spine. Mean standardized uptake values (SUVs, normalized to muscle) were calculated for each ROI. Subsequently, ^89^Zr–MS-Hu6 uptake of each tissue was expressed as the average of all mean SUV values *per* organ. After imaging, the mice were sacrificed and perfused with 20 mL of PBS and tissues of interest, namely brain, heart, kidney, pancreas, liver, lung, bone, bone marrow, BAT, subcutaneous WAT, visceral WAT, adrenal, blood, testis, spleen, and muscle, were isolated for γ–counting..

For biodistribution studies using AF750–labelled MS-Hu6, we first imaged the whole body 16 hours after injection using IVIS Spectrum *In Vivo* Imaging System (Perkin Elmer). Mice were then perfused with PBS, sacrificed and organs, namely heart, thymus, brain, gastronemus and soleus muscle, BAT, adrenals, liver, gall bladder, spleen, kindey, subcutaneous WAT, visceral WAT gonadal WAT, testes and bone were removed and imaged using the same IVIS platform to calculate average radiance efficiency *per* square area.

### Biodistribution and Safety Studies in Monkeys

After an overnight fast, two male *Cynomolgus* monkeys [Scott, aged 14 years, weight 9.8 kg; and Andy, aged 15 years, weight 6.15 kg] were anesthetized with ketamine (5.0 mg/kg) and dexemedetomidine (0.0075 to 0.0015 mg/kg). The monkeys were injected with ^89^Zr–MS-Hu6, and blood was drawn at 30 minutes, and at 48 and 120 hours from the tail vein. Vitals, including mean arterial, systolic and diastolic blood pressure, respiratory rate, heart rate and rectal temperature, were recorded using Waveline Touch system (DRE) and Welch Allyn rectal thermometer. PET and MR images were acquired on a combined 3T PET/MRI system (Biograph mMR, Siemens Healthineers). Whole body MR images from each PET bed (head, thorax, pelvis) were automatically collated together with a scanner. MR parameters were as follows: acquisition plane, coronal; repetition time, 1000 ms; echo time, 79 ms; number of slices, 224; number of average, 2; spatial resolution of 0.6 mm x 0.6 mm x 1.0 mm; and acquisition duration, 29 min and 56 s *per* bed. After acquisition, PET raw data from each bed were reconstructed and collated offline using the Siemens proprietary e7tools with an ordered subset expectation maximization (OSEM) algorithm with point spread function (PSF) correction. A dual–compartment (soft tissue and air) attenuation map was used for attenuation. Image analysis was performed using Osirix MD, version 11.0. Whole-body MR images were fused with PET images and analyzed in an axial plane. Regions of interest (ROIs) were drawn on various tissues. The liver, kidney, BAT (interscapular region), subcutaneous WAT, visceral WAT, gonadal WAT, gallbladder, spleen, brain, and testes were traced in their entirety; bone marrow was imaged from the shoulder; and 3 lumbar vertebrae; and muscle was imaged from the quadriceps. Mean SUVs were calculated for each ROI. ^89^Zr–MS-Hu6 uptake of each tissue was expressed as the average of all mean SUV values *per* organ. Serum was collected for blood chemistry analysis by IDEXX BioAnalytics.

### ELISPOT Assay

Human peripheral blood mononuclear cells (PMBCs, obtained from Immunospot, Cellular Technology Ltd.) were cultured for 12 days in FBS–free DMEM with regular medium change, and plated at a density of 10^5^ cells/well in ImmunoSpot ELISPOT plates. Cells were then exposed to MS-Hu6, CEFT or DMEM for 48 hours, following which IL-2 and IFNγ expressing cells were quantitated *per* manufacturer’s instructions.

### Target Engagement Assays

For pharmacodynamic studies—namely engagement of MS-Hu6 with FSH as its target— male and female C57BL/6 mice were injected i.p. with MS-Hu6 or human IgG (200 µg, for each). After 16 hours, mice were bled and total IgG (mouse and human) was pulled down using protein A beads (ThermoFisher, Catalog # 20333). An in–house sandwich ELISA was used to measure human IgG (control IgG or MS-Hu6) in the eluate. For this, plates were coated with anti–human Fc (Sigma, Catalog # FAB3700259) and goat anti–human HRP–conjugated IgG (H+L) (Invitrogen, Catalog # A18805) was used to for capture total human IgG. The same elute then underwent a second in–house sandwich ELISA, in which the plate was coated with anti–human Fc (Sigma, Catalog # FAB3700259), and after overnight incubation at 4°C, the MS-Hu6–FSHβα complex was captured by an antibody to the α subunit of FSH. In a separate study, a commercial ELISA for FSH (Biotechnology Systems, Catalog # M7581) was used to determine whether MS-Hu6 binding to FSH interfered with its detection.

### *In Silico* Analyses

We used Protein–Sol, a computational algorithm based on machine learning of amino acid sequences and physicochemical variables from 137 antibodies that are FDA–approved or in late– stage clinical development (https://protein-sol.manchester.ac.uk/abpred) (32–34). Protein–Sol uses antibody sequences (V_H_ and V_L_) as inputs to provide predicted outputs for clone self–interaction by bio–layer interferometry (CSI-BLI), poly–specificity reagent (PSR), baculovirus particle ELISA (BVP-EL), cross–interaction chromatography (CIC), ELISA, accelerated stability (AS), hydrophobic interaction chromatography (HIC), stand–up monolayer adsorption chromatography (SMAC), salt gradient affinity capture (SGAC), expression titer in HEK cells (HEK), affinity capture (AC), and differential scanning fluorescence (DSF). It also provides a meta value for each sequence (1=best; 100=worst) by averaging ranks for 8 experimental parameters. For predicting the pI value MS-Hu6, we inputted its sequence into Expasy (Swiss Bioinformatics Research Portal, https://web.expasy.org/compute_pi/). For predicting the “humanness” of humanized MS-Hu6 *versus* parent chimera and other humanized or fully human molecules (26), we inputted sequences into abYsis (version 3.4.1; http://www.abysis.org/abysis/sequence_input/key_annotation/key_annotation.cgi) to obtain *Z*-scores, and compared the scores with that of V_H_ and V_L_ĸ chain of IgG1.

### Protein Thermal Shift Assay

The thermal shift assay used a fluorescent reporter, Sypro–Orange (Protein Thermal Shift Dye Kit, ThermoFisher, Catalog # 4461146), to detect hydrophobic domains that are exposed following the heat–induced unfolding of globular proteins. MS-Hu6 (1.5 µg/µL), formulated or in PBS, was incubated with or without human FSH (0.5 µg/µL) at room temperature for 30 minutes, with fluorescence captured sequentially at 0.3 °C increments using a StepOne Plus Thermocycler (Applied Biosystems). T_m_ was calculated based on the inflection point of the melt curve, and thermal shift was derived from ΔT_m_ = T_m_A–T_m_B.

### Isoelectric Focusing

For determining the isoelectric pH (pI), two dimensional electrophoresis was performed by first rehydrating MS-Hu6 (500 µg) for 2 hours at room temperature in rehydration buffer (8M urea, 2% CHAPS, 0.5% IPG buffer, and trace of bromophenol blue) without DTT. The sample was then run on an 18 cm 3-10 strip using the Ettan IPGphor 3 Isoelectric Focusing System (GE Healthcare). Four voltage steps (50 V for 10 hours; 500 V for 1 hour; 1000 V for 1 hour; 8000V for 4 hours) were followed by Coomassie blue staining.

### Chromatography

An NHS ester column (HiTrap NHS Activated HP, Cytiva, Catalog # 17071601) was conjugated with either human IgG (for CIC) or MS-Hu6 (for SIC) in 0.2 M NaHCO_3_, 0.5 M NaCl (pH 8), *per* manufacturer. Unbound IgG was removed by successive washings (x3) at a rate of 0.4 mL/minute using Buffer A (0.5 M ethanolamine, 0.5 M NaCl, pH 8,3) and Buffer B (0.1 M sodium acetate, 0.5 M NaCl, pH 4). An unconjugated column was prepared without including any IgG in the coupling step. MS-Hu6, either as a formulation or in PBS, was run through the unconjugated column, followed by either conjugated columns at 0.2 mL/min. For SEC, we used a Superdex 200 Increase 10/300 GL (GE Lifesciences, Catalog # 28990944) and passed MS-Hu6 (2 mg) in either PBS or formulation buffer after 3 cycles of freeze–thaw or incubation for 1 week at 4 °C, 37 °C or 50 °C. For HIC, we used HiTrap Butyl FF Sepharose column (GE Lifesciences, Catalog # GE17-1357-01) to run MS-Hu6 at 1 mL/minute in a linear gradient from 1 M to 0 M (NH_4_)_2_SO_4_ [generated using 1.8 M (NH_4_)_2_SO_4_ and 0.1 M Na_2_PO_4_ (pH 6.5), and 0.1 M Na_2_PO_4_ (pH 6.5)]. For CIC, SIC, SEC, HIC, absorbance was monitored at 280 nm using the AKTA PURE FPLC (GE Lifesciences), and the data analyzed by Unicorn version 6.4.

### Polyspecificity Testing

We adapted a previous method to antibody determine polyspecificity (34), and developed an in–house ELISA by coating plates with multiple antigens (50 nM for each), namely double stranded DNA (Shear Salmon Sperm DNA, 5’-3’ Inc., Catalog # 5302-754688), single stranded DNA (Deoxyribonucleic acid, single stranded from Calf Thymus, Sigma, Catalog # D8899), cardiolipin (Cardiolipin sodium salt from bovine heart, Sigma, Catalog # C0563), LPS-B5 (Lipopolysaccharide *E*. *coli* 055:B5, Calbiochem, Catalog # 437625,), LPS-B4 (LPS-EB, InvivoGen, Catalog # tlrl-eblps, 50 nM), KLH (keyhole limpet, Hemocyanin from *Megathura crenulata*, Sigma, Catalog # H8283), insulin (Humulin R, Lilly, Catalog # HI213), baculovirus particle (Medna, Catalog #: E3001), flagellin (Adipogen, Catalog # AG-4013-0095-C100), human serum albumin (Sigma, Catalog # A9511) and human pituitary FSH (National Hormone and Pituitary Program, UCLA). The antigens were exposed MS-Hu6 (100 nM), overnight at 4 °C, and any antigen–MS-Hu6 complex was captured by goat anti–human HRP–conjugated IgG (Invitrogen, Catalog # A18805).

### Statistical Methods

Statistically significant differences between any two groups were examined using a two– tailed Student’s *t*-test, given equal variance. *P* values were considered significant at or below 0.05. Data points were excluded if they were 2.5 standard deviations above or below means

#### Ensuring Rigor and Reproducibility

There is a nascent movement to ensure that preclinical data is true and accurate (58–63). M.Z. and C.J.R. coined the phrase ‘contemporaneous reproducibility,’ which refers to the synchronous reproduction of data in more than one laboratory. As Zaidi’s discovery of the effects of FSH on body fat were novel and unexpected, he reached out to C.J.R. for help in form of a reproducibility study. Key data sets were reproduced by C.J.R in a process that lasted over three years, as other validation studies were added by both laboratories. The term replicability refers to the ability of one or more independent groups to replicate a finding using a different technology or method––replicability is a measure of truth or significance of a given finding(64). Here, we have replicated a key finding that Hu6 increased bone mass in the M.Z. and C.J.R. labs, with µCT data independently produced by J.C. To further enhance transparency, we have hosted detailed procedures and raw datasets on our GLP– compliant MediaLab Document Control System that all investigators have access to. All data have undergone quality checks before the final product was signed off. Such practices requiring unfettered transparency remain fundamental to ensuring rigor.

## LEGENDS TO FIGURES

**Supplementary Figure 1:**
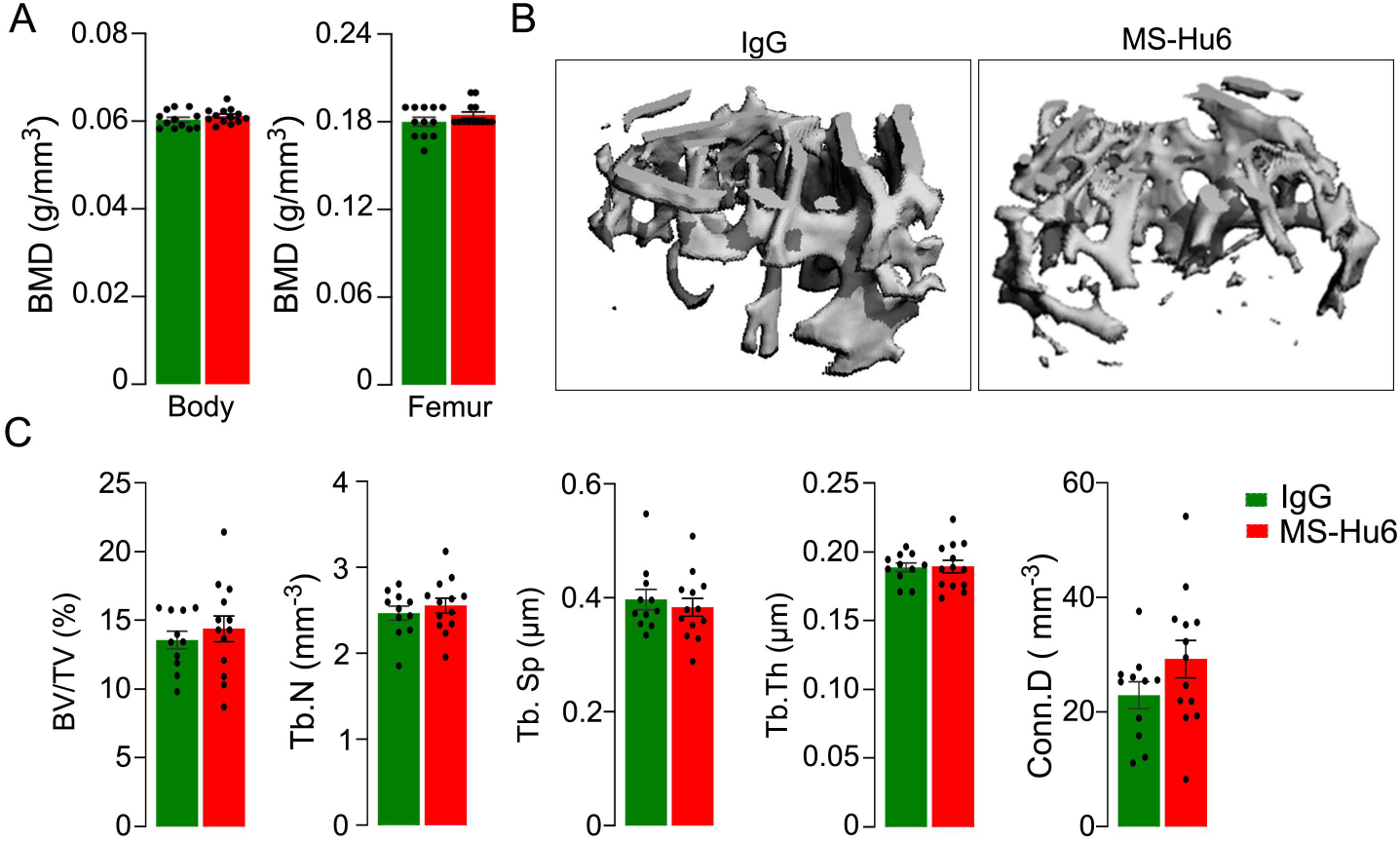
Bone Phenotype of the C3H Mouse is Resistant to Anabolic Actions of MS-Hu6. *Piximus* measurements of total body and femur bone mineral density (BMD) **(A),** as well as µCT images **(B)** and quantitative estimates of fractional bone volume (BV/TV), trabecular number (Tb.N), spacing (Tb.S.) and thickness (Tb.Th), and connectivity density (Conn.D) (C). Experiment performed at C.J.R. lab, with µCT analysis independently at J.C’s lab.

